# Temporal Dynamics of Sensorimotor Integration for Auditory and Proprioceptive Target Information During Movement Preparation and Execution

**DOI:** 10.1101/2025.07.09.663821

**Authors:** Ivan Camponogara

## Abstract

Endpoint movement variability in non-visual (auditory and proprioceptive) target reaching is modality-dependent. This study investigated whether such modality-specific effects may emerge during movement preparation and how they unfold during movement execution. Participants performed reaching movements toward auditory, proprioceptive, or audio-proprioceptive targets. Electromyography and movement kinematics were recorded to examine the effect of target-related sensory modality on sensory encoding and coordination of motor commands during movement preparation, as well as their effect on movement execution. Results revealed a modality-dependent sensory encoding phase, whereas a modality-independent coordination of motor commands. While the target modality did not affect movement initiation, it modulated the subsequent movement execution. These findings demonstrate that, in non-visual target reaching, the effect of target-related sensory modality extends beyond execution to specific phases of movement preparation. Our results support a four-stage model of action control toward non-visual targets, consisting of a modality-dependent sensory encoding phase of movement preparation and early execution, as well as a modality-independent motor coordination and movement initia-tion. These findings provide new insights into the temporal dynamics of sensorimotor control without vision, suggesting a different use of non-visual (proprioceptive and auditory) target-related sensory information during distinct phases of movement preparation and execution.

**Highlights:**

- Auditory and proprioceptive target reaching shows modality-specific endpoint variability.
- This study investigates whether such modulation occurs within the movement preparation stage.
- Modality-specific sensory modulation has been found in specific phases of movement preparation.
- Modality-specific effects emerge during early and late movement execution.
- Findings support a four-stage model of non-visual action control, separating encoding, coordination, initiation, and execution.

## Introduction

In everyday life, we often perform actions toward targets that we can’t see, leveraging information coming from auditory and proprioceptive sensory modalities. For instance, we mostly use audition to catch a flying mosquito in the dark, while proprioception guides us if the mosquito lands on our arm. Studies have shown that, even though actions toward *non-visual* targets (auditory and proprioceptive) subserve the same goal (successfully reaching a target), they exhibit modality-specific patterns of variability, with actions toward auditory targets exhibiting greater variability than those toward proprioceptive targets. Concurrently, actions toward proprio-ceptive targets are similar to those toward audio-proprioceptive targets, suggesting a reliance on proprioception in multisensory conditions (Camponogara, 2025). However, while providing evidence on the role of *non-visual* target-related information on action performance, these findings are grounded in studies examining endpoint accuracy and precision (Cuppone et al, 2018, 2019; Camponogara, 2025). As a result, current evidence offers insight into how different target-related sensory information is used *during* movement execution, when refining ongoing actions (Elliott et al, 2016; Brenner and Smeets, 2025). This raises the critical question of whether differences in auditory and proprioceptive target reaching are already evident *before* movement onset, during movement preparation.

Studies on visual and proprioceptive/haptic target reaching suggest that the modality-dependent modulation of action performance emerges already during movement preparation, prior to movement onset (Camponogara, 2023). Visual exposure to a haptically sensed target before movement onset significantly reduces reaction time (RT) (Camponogara et al, 2024), the index of movement preparation (Haith et al, 2016). Even brief visual exposure to a haptic target during the RT interval improves the subsequent movement performance compared to purely haptic or proprioceptive reaching (Desmurget et al, 1997; Monaco et al, 2010; Jones et al, 2012; Camponogara and Volcic, 2021), further underlining the fundamental role of sensory inputs in movement preparation. Additional evidence for modality-specific preparation is provided by studies reporting distinct movement trajectories (Sarlegna et al, 2009) and visuomotor-adaptation effects (Sober and Sabes, 2003, 2005; Bernier et al, 2007) toward visual versus proprioceptive targets approximately ∼ 70 ms after movement onset, highlighting different sensorimotor transformation processes engaged during the preparation of visually versus proprioceptively guided actions. Neuroimaging and electrophysiological findings converge with these behavioral results, revealing modality-specific activation patterns in visual and somatosensory cortical areas before the movement onset (Blouin et al, 2014; Manson et al, 2019; Bianco et al, 2020b; Kreter et al, 2025) and different temporal patterns of motor-related cortical areas when reaching visual and proprioceptive targets (Bernier et al, 2009). Thus, taken together, these studies suggest that movement preparation is contingent on the target-related sensory modality.

It is, however, worth considering that such evidences are mainly based on studies investigating actions toward *visual* and *non-visual* (i.e., proprioceptive) targets, which may not generalize to sensory modalities that are exclusively *non-visual*, such as proprioception and audition. A subtle yet crucial difference characterizes visual vs. proprioceptive and auditory vs. proprioceptive target reaching, which concerns the amount of sensory modalities monitoring the *reaching hand* throughout the movement. In visual target reaching, the *reaching hand* is sensed by both vision and proprioception, allowing a continuous multisensory monitoring of its position from movement preparation to completion. In contrast, in proprioceptive and auditory target reaching, the position of the *reaching hand* is monitored solely through unisensory proprioceptive inputs. Previous studies have shown that the availability of multisensory information in visual target reaching accounts for the differences in movement preparation observed when actions toward visual and proprioceptive targets are compared. As a matter of fact, removing or altering visual information about the *reaching hand* has a substantial impact on movement preparation toward a visual target (Sarlegna and Sainburg, 2007; Sober and Sabes, 2005; Sarlegna et al, 2009). Thus, the different movement preparation usually reported in previous studies on visual vs. proprioceptive target reaching may not generalize to auditory and proprioceptive target reaching, where the *reaching hand* is monitored through proprioceptive modality only, and the target is defined via either audition or proprioception.

In this regard, it is unclear whether reaching movements toward auditory and proprioceptive targets rely on similar or modality-specific movement preparation. Behavioral studies have shown different movement preparation for actions toward proprioceptive targets when cued by an auditory or a somatosensory stimulus, with higher RT in the auditory compared to the somatosensory condition (Manson et al, 2019). Concurrently, EEG studies have shown different predictive patterns of somatosensory and auditory cortex activity before stimulus presentation in both sensory discrimination and motor tasks (Bianco et al, 2020b,a). Together, these studies suggest a modality-specific movement preparation for actions directed toward non-visual targets. In contrast, previous fMRI studies have found that auditory and somatosensory inputs are processed within an integrated neural system (Farn’e and Làdavas, 2002), which includes both sensory (Kayser et al, 2005; Caetano and Jousmäki, 2006; Zierul et al, 2017; Lohse et al, 2022) and premotor brain areas (Serino, 2019; Camponogara, 2023); also involved in action preparation (Camponogara, 2023). Moreover, studies reported comparable prefrontal and motor cortex activation following auditory and somatosensory target presentations (Bianco et al, 2020a; Fiorini et al, 2021; Lucia et al, 2023). Combined with previous studies on non-visual target reaching (Camponogara, 2025), these studies suggest that movement preparation might be similar across non-visual modalities, and the target-related sensory modality may be mainly relevant after the movement onset, during the movement execution. Thus, building on these findings, two main hypotheses emerge. First, movement preparation and execution may be both modality-specific, with the former exhibiting distinct characteristics according to the target-related sensory modality. Alternatively, actions directed toward auditory and proprioceptive targets may exhibit similar movement preparation and early execution, with sensory modality becoming particularly relevant in the later stages of movement for fine-tuning and adjustment (Elliott et al, 2016; Brenner and Smeets, 2025). The present study aims to shed light on how movements toward *non-visual* (auditory and proprioceptive) targets may unfold during their preparation and execution stages.

The sensorimotor integration process during movement preparation was assessed through electromyography (EMG) recordings and kinematic analysis. These measures were used to dissociate two temporal components of movement preparation, measured through the reaction time (RT): the premotor reaction time (PRT) and motor reaction time (MRT). Each component characterizes a specific sensorimotor process within movement preparation. The PRT is the time from the stimulus onset to the first visible muscle activation. It corresponds to the time taken for the central nervous system to encode the sensory stimulus. The PRT is considered the central component of motor preparation (Ballanger and Boulinguez, 2009), which is the process by which the central nervous system integrates sensory inputs with motor commands. The MRT, instead, is the time from the first muscle activation to the onset of movement kinematics and corresponds to the timing of coordinating the motor response. This involves defining the muscle activation from alpha motor neurons to initiate movement and executing the actual action response (Weiss, 1965; Botwinick and Thompson, 1966; Ballanger and Boulinguez, 2009). The MRT represents the peripheral component of motor preparation (Ballanger and Boulinguez, 2009), which is related to the process coordinating the motor commands (Schmidt and Alan Stull, 1970; Schmidt et al, 1979; Harris and Wolpert, 1998). Both phases are affected by stimulus characteristics and task demands (Botwinick and Thompson, 1966; Schmidt and Alan Stull, 1970; Schmidt et al, 1979; Van Donkelaar and Franks, 1991; Nikravan et al, 2021; Servant et al, 2021) and are independent: a change in PRT does not correspond to a change in MRT (Botwinick and Thompson, 1966; Schmidt and Alan Stull, 1970).

Thus, while investigating RT allowed us to determine whether the sensory modality generally affects the motor preparation phase, studying each specific component of action preparation enabled us to define whether its mod-ulation is related to the sensory encoding process or the coordination of motor commands. Action preparation was further investigated by analyzing the EMG activity during the MRT, the peak acceleration, the hand posi-tion, and its variability in depth and azimuth directions at the peak acceleration. Peak acceleration was selected because it provides insight into the sensorimotor processes involved in action preparation (Sarlegna and Sain-burg, 2007, 2009). The role of each sensory modality in the action execution phase, instead, was investigated by analyzing peak velocity, peak deceleration, movement time, as well as the reaching finger position and variability in both depth and azimuth directions at the peak velocity, peak deceleration, and the end of the movement. If the target-related sensory modality affects action performance already during the motor preparation phase, actions toward auditory and proprioceptive targets will exhibit different preparation timings (RT, PRT, MRT), EMG activity, and different movement kinematics at immediate proximity of movement onset (i.e., at the peak acceleration). In contrast, if target-related sensory information is fundamental only at the late stage of the movement, neither movement preparation (RT, PRT, MRT, EMG activity) nor initial action execution will be affected by the target modality. In this case, we expect a different pattern of movement kinematics during early and late movement execution only (PV, Dec, MT, and finger positions at these kinematic markers).

## Material and Method

### Participants

We tested a total of thirty-nine participants. However, eight participants were excluded from the final analysis due to the high background noise in the electromyographic data, which prevented defining the onset of muscle activation. Thus, the final sample included thirty-one participants (right-handed, 11 males, age 32.8 ± 13.8 years). All participants self-reported normal hearing and no known history of neurological disorders. All of the participants were naïve to the purpose of the experiment. The experiment was undertaken with the understanding and informed written consent of each participant, and the experimental procedures were approved by the Institutional Review Board of Zayed University, Abu Dhabi.

### Apparatus

The custom-made experimental apparatus consisted of a perforated wooden board of 600 by 600 mm supported by four 100 mm high pedestals placed on the experimental table. The board was perforated throughout its entire surface to avoid participants using the holes as tactile feedback when landing on the speaker’s position. Three speakers (Gikfunk, 4O 0hm 3 Watt, 40 mm diameter, 30 mm height) were placed underneath the board, powered through a USB cable, and connected with an external soundcard (Motu 4Pre) via a mini amplifier (PAM8403). The center of the speakers was placed 200 mm far at 0 (i.e., in front), 45, and 90 degrees on the left with respect to a 5 mm high rubber bump that acted as a starting point (Figure 1 A). The bump was placed 100 mm far from the side of the wooden board closest to the subject position. A 5 mm high rubber bump was placed underneath the center of each speaker and was used as a reference for the left index position during the proprioceptive and audio-proprioceptive conditions. The right-hand index finger movement was acquired online at 200 Hz with sub-millimeter resolution using an OptiTrack system (NaturalPoint, Inc.). The muscle activation of the Anterior Deltoid was recorded using a surface electromyography (EMG) system (four-channel Wave Plus EMG system Mini, Cometa Srl). This muscle was chosen because it is the first muscle to activate when performing an arm flexion (Latash et al, 1995). A chin rest was placed on the front side of the wooden board to avoid head movements during the experiment. The chin position was set at 25 cm height from the wooden surface, and the chair height was adjusted at the participants’ comfortable height. A pink noise of 1000 ms duration (65 db) was used as the auditory target. The same noise was used to signal the start of the trial in the proprioceptive condition, but delivered from external speakers. A 100 ms duration pure tone of 500 Hz (65 dB) was used to signal the end of the trial. Both the pink noise in the proprioceptive condition and the pure tones (signaling the end of the trial) were delivered through two external speakers (Logitech 980-000816 Z150). The motion capture, electromyography, and sound delivery were controlled via MATLAB (MathWorks).

**Figure 1:**
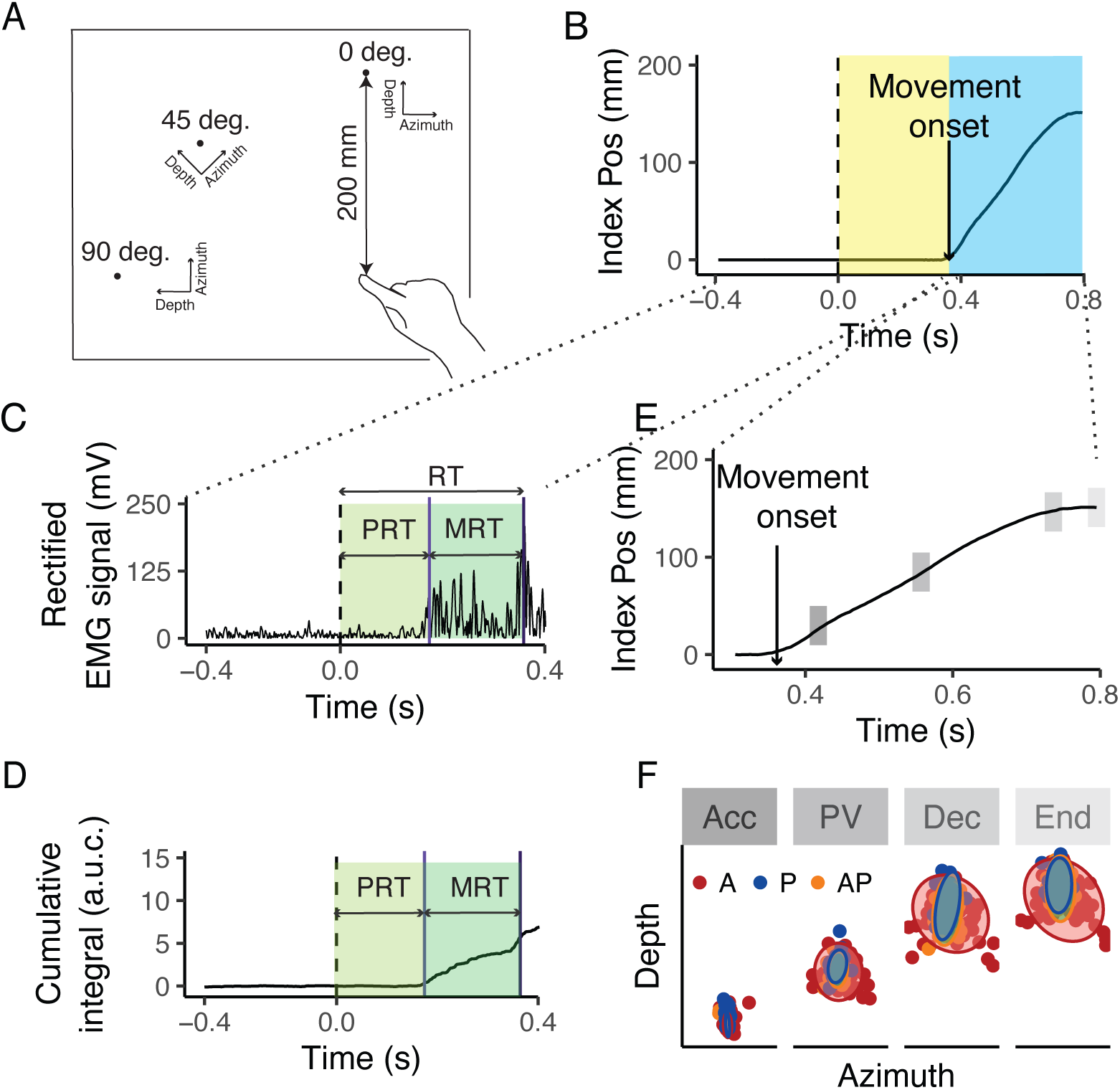
Experimental setup and signal analysis. A) Top view of the experimental setup. Targets were positioned at 200 mm in front (0 degrees) or laterally (45 and 90 degrees) from a start position. B) Index positional signal along the depth dimension. The yellow area represents the movement preparation phase (i.e., Reaction Time), whereas the blue part represents the movement execution phase. The 0 in the time axis represents the time of auditory stimulus presentation (t0). C) Band-pass filtered and rectified EMG signal, and D) Integrated (cumsum function) EMG signal. The dashed black line represents the t0, whereas the light blue and dark blue lines represent the time of muscle burst and movement onset, respectively. The light and dark green areas represent the premotor reaction time (PRT) and motor reaction time (MRT). E) Index position along the depth dimension. The shaded grey areas represent the positions at which the maximum acceleration, peak velocity, and deceleration occurred, as well as the endpoint position. These are the positions along the trajectory where movement position and variability were assessed. F) Index finger positions (dots) in azimuth and depth, and their 95% confidence ellipses of a representative participant. Each panel represents the finger position at the Acc, PV, Dec, and Endpoint.

### Procedure

Participants were blindfolded for the whole duration of the experiment. All the trials started with the participant’s head on the chin rest, the index digit of the right hand positioned on the start position, and the left hand on the left side of the chin rest. Two splints were used to immobilize the wrist and index finger of the reaching hand in a fixed position, allowing arm movements to occur only through shoulder and elbow rotation (van Beers et al, 2004). In an Auditory condition (A), the pink noise was delivered from the target speaker, and participants had to reach the point where they perceived the noise coming from. In a Proprioceptive (P) and Audio-Proprioceptive (AP) condition, before each trial, participants were instructed to move the index finger of the left hand underneath the specified target speaker over the bump placed at the center of it. As soon as the finger was placed on the bump, in the P condition, the pink noise was delivered from the external speakers, whereas in AP, the pink noise was delivered from the speaker over the left finger. As soon as participants heard the pink noise, they had to reach the point where they thought the tip of their left index finger was, or both the tip of the finger and the pink noise were delivered by raising the finger from the start position. In all sensory conditions, after 2 seconds from the onset of the pink noise, the end tone was delivered, and participants had to return their right hand to the start position and their left hand to the side of the chin rest in the P and AP conditions. Before each experiment, participants underwent a practice phase in the A condition, consisting of 10 trials, during which they became accustomed to the task. We performed 20 trials for each target, leading to 60 trials per condition (180 trials per participant). Conditions were randomized between participants.

### Data analysis

Kinematic and EMG data were analyzed in R (R Core Team, 2020). For the kinematic data, the x and y coordinates of the index and speakers’ positions were smoothed and differentiated with a third-order Savitzky-Golay filter with a window size of 21 points. These filtered data were then used to compute velocities and accelerations of the Index finger in two-dimensional space. Movement onset was defined as the moment of the lowest, non-repeating Index finger acceleration value before the continuously increasing Index finger acceleration values, while the end of the movement was defined by applying the same algorithm but by reversing the signal (Camponogara and Volcic, 2021). Trials in which the movement onset or end was not correctly detected were discarded from the analysis (310 trials, 6.2% of the total number of trials). From the kinematic data, we computed the Reaction Time (RT), defined as the time from the stimulus to the movement onset. Additionally, we computed the maximum index finger Acceleration (Acc), Peak Velocity (PV), Deceleration (Dec), and movement time (MT). Movement position along the depth and azimuth direction was also extracted at each kinematic marker (Acc, PV, Dec) and at the endpoint (Figure 1E, F). Then, we calculated the distance of the finger from the target along both directions at Acc, PV, Dec, and Endpoint. Such distance was the object of the final analysis. The kinematic signals were also resampled in 201 points evenly spaced along the three-dimensional trajectory in the range from 0 (movement onset) to 1 (movement end) in 0.005 steps using cubic spline interpolation (Camponogara and Volcic, 2019). Such re-sampling was used to define the point along the finger trajectory where the peak acceleration (pACC), peak velocity (pPV), and peak deceleration (pDec) occurred, expressed as a proportion of the total index finger trajectory at the specific kinematic marker.

From the EMG signal, we computed the Premotor Reaction Time (PRT) and the Motor Reaction Time (MRT). These were computed by first detecting the onset of the muscle burst. The EMG data were first aligned with the kinematic data (Figure 1B, C), then band-pass filtered at 20-450 Hz with a second-order Butterworth filter and rectified. For each signal, the baseline muscle activity was then calculated as the mean muscle activity from the start to 300 ms following the beginning of the trial. This was subtracted from the EMG signal (Contemori et al, 2022). We then calculated the cumulative integral of the signal using the *cumtrapz* function in R. The onset of muscle activation was calculated by considering the time the cumulative integral (Servant et al, 2021) exceeded a threshold of 0.5 (Figure 1D). The accuracy of onset detection was ensured by a trial-by-trial visual inspection of the EMG signal, which confirmed a more accurate detection of the muscle onset than the classic use of baseline and standard deviation approach in rectified low-pass EMG signals (Bertucco and Cesari, 2010; Delmas et al, 2018). The method of cumulative integral was of particular benefit for signals with a relatively high baseline value. Trials in which the muscle background activity was too high and prevented the automatic detection of the muscle onset were discarded (501 trials, 9% of the total number of trials). We then extracted the PRT, defined as the time from the sound presentation to the onset of muscle burst, and the MRT, defined as the time from the onset of muscle burst to the movement onset. The EMG signal was further processed to quantify muscle activation during the MRT. The integral of the EMG signal was computed within the MRT interval using the *trapz* function in R. The average baseline activity was obtained over a 300 ms window ranging from the start of the trial to 300 ms afterward. The integrated EMG values during MRT were then baseline-corrected. For each subject, the EMG activation was then normalized to the maximum EMG amplitude observed across all trials within each condition. As a result, the final normalized EMG values ranged from 1 to -1 (Bertucco et al, 2021).

#### Statistical analysis

The MT, RT, PRT, MRT, pAcc, pPV, and pDec were modeled through Bayesian linear mixed-effects regression models. The EMG at MRT, the index finger position at Acc, PV, Dec, and the end of the movement along the azimuth and depth directions were modeled through Bayesian distributional linear mixed-effects regression models. All models were estimated using the brms package (Bürkner, 2017), which implements Bayesian multilevel models in R using the probabilistic programming language Stan (Carpenter et al, 2017). To avoid models’ divergencies, we excluded from the analysis trials where RT, PRT, and MRT were lower than zero (i.e., participants anticipated the response or muscle burst occurred at the same time as movement onset). These corresponded to 93 trials in total (1.6% of the data). All the models included as predictors the categorical variable Condition (A, P, and AP) and the continuous variable Position (0, 45, and 90 degrees), allowing for the estimation of the average for each dependent variable and its change as a function of the target position (slope). All the models included the independent random (group-level) effects for subjects.

We fitted separate models for each timing (MT, RT, PRT, MRT) and proportion of trajectory (pACC, pPV, pDec) variable. The timing variables (MT, RT, PRT, MRT) were fitted using a shifted log-normal distribution (Bürkner, 2017). The model included weakly informative prior distributions for each parameter to provide information about their plausible scale. We used normally distributed priors for the Condition and Position fixed-effect predictor (*β_Condition_*: mean = 5 and sd = 4; *β_Position_*: mean = 0 and sd = 1), whereas for the group-level standard deviation parameters we used zero-centered Student *t* -distribution priors, with scale values defined by the “get prior” function. (*df* = 3, scale = 2.5). Finally, we set a prior over the correlation and residual correlation matrix that assumes that smaller correlations are slightly more likely than larger ones (LKJ prior set to 2). Using a shifted log-normal distribution, the model returned a log-transformed posterior distribution of intercepts and slope. Thus, the intercept was back-transformed to the original scale by exponentiation, whereas the slope was expressed as a percentage change of MT, RT, PRT, and MRT for each change in degree of target position (Greenwood, 2022).

For the pAcc, pPV, and pDec, we used a zero-one inflated beta distribution (Bürkner, 2017), as variables were proportions (i.e., percentage of space covered at peak acceleration, velocity, and deceleration). The model was fitted considering weakly informative prior distributions for each parameter to provide information about their plausible scale. We used normally distributed priors for the Condition and Position fixed-effect predictor (*β_Condition_*: mean = 0 and sd = 1; *β_Position_*: mean = 0 and sd = 0.05), whereas for the group-level standard deviation parameters we used zero-centered Student *t* -distribution priors, with scale values defined by the “get prior” function. (*df* = 3, scale = 2.5). Finally, we set a prior over the correlation and residual correlation matrix that assumes that smaller correlations are slightly more likely than larger ones (LKJ prior set to 2).

For the EMG at MRT, Acc, PV, and Dec, we fitted separate distributional regression models, which allowed us to estimate their posterior distribution and standard deviation SD (sigma), taking into account both within and between-subject variability. We used a Gaussian distribution for the estimation of the variables. The model was fitted considering weakly informative prior distributions for each parameter to provide information about their plausible scale. We used normally distributed priors for the Condition and Position fixed-effect predictor (*β_Condition_*EMG mean = 0, sd = 1, Acc mean = 8000, sd = 500; PV mean = 700, sd = 100; Dec mean = -4000, sd = 500 *β_Position_*: EMG mean = 0, sd = 0.05, otherwise mean = 0 and sd = 10 for all the other variables), whereas for the group-level standard deviation parameters we used zero-centered Student *t* -distribution priors, with scale values defined by the “get prior” function. (*df* = 3, EMG scale = 2.5, Acc scale = 3210.6, PV scale = 196, Dec scale = 1670). Finally, we set a prior over the correlation matrix that assumes that smaller correlations are slightly more likely than larger ones (LKJ prior set to 2).

Separate distributional regression models were also used to fit the index finger positions in azimuth and depth directions at Acc, Dec, PV, and Endpoint. The model allowed us to estimate the posterior distribution of the index position and SD (sigma) in both depth and azimuth directions, taking into account both within and between-subject variability. The models included the index position in azimuth and depth as the dependent variable, and Condition and Position as predictors. Thus, the models estimated the average position in the depth and azimuth directions, and the average SD in the depth and azimuth directions. The models included the independent random (group-level) effects for subjects. The model was fitted considering weakly informative prior distributions for each parameter to provide information about their plausible scale. We used normally distributed priors for the Condition fixed-effect predictors (*β_Condition_* in depth: Acc mean = -200 and sd = 50; PV mean = -100 and sd = 30; Dec mean = -10 and sd = 50; End mean = 0, sd = 50, in azimuth mean = 0 and sd = 10 for all the variables) *σ_Condition_* in azimuth and depth: mean = 1 and sd = 1). For the group-level standard deviation parameters, we used zero-centered Student *t* -distribution priors, with scale values defined by the “get prior” function. (Acc azimuth = *df* = 3, scale = 4.3, depth = *df* = 3, scale = 5.2; PV azimuth = *df* = 3, scale = 15.8, depth = *df* = 3, scale = 31.1; Dec azimuth = *df* = 3, scale = 29.9, depth = *df* = 3, scale = 48.8; End = azimuth = *df* = 3, scale = 32.6, depth = *df* = 3, scale = 38.9). Finally, we set a prior over the correlation and residual correlation matrix, assuming that smaller correlations are slightly more likely than larger ones (LKJ prior set to 2).

For all models, we ran four Markov chains simultaneously, each for 6,000 iterations (1,000 warm-up samples to tune the MCMC sampler) with the delta parameter set to 0.99, resulting in a total of 20,000 post-warm-up samples. Chain convergence was assessed using the *R̂* statistic (all values equal to 1), visual inspection of the chain traces, and a bulk effective sample size higher than 1000 (Bürkner, 2017).

#### Statistical Inference

The posterior distributions represent the probabilities of the parameters conditional on the priors, model, and data, and they represent our belief that the “true” parameter lies within some interval with a given probability. We summarize these posterior distributions by computing the medians and the 95% Highest Density Intervals (HDI). The 95% HDI specifies the interval that includes, with a 95% probability, the true value of a specific parameter. To evaluate the differences between the two compared conditions, we have subtracted the posterior distributions between specific conditions. The resulting distributions are denoted as the credible difference distributions and are again summarized by computing the medians and the 95% HDIs. For statistical inferences, we assessed the overlap of the 95% HDI with zero. A 95% HDI of the credible difference distribution that is not included within the zero region is taken as evidence that the two conditions differ from each other for that specific variable. For the sake of clarity, from now on, we will refer to the SD results as variability.

## Results

### Movement Preparation

#### Effect of the sensory condition

Results showed that reaching an auditory target requires a longer RT and PRT compared to proprioceptive and audio-proprioceptive targets (Figure 2, panels from A to D). Concurrently, the analysis revealed a slightly higher MRT in A compared to P and AP conditions (Figure 2 E, F). However, the lower boundary of HDI differences was very close to zero (∼0.1 ms), which prevents drawing strong conclusions regarding differences between non-visual conditions. The normalized EMG activity and its variability was similar across conditions (Figure 2 G, H), except for a slightly higher variability in A compared to AP (Figure 2 I, J). The Acc and its variability were similar across conditions (Figure 3A, B), as well as the pAcc, which occurred at the ∼5 % of the movement trajectory (Figure 4 A, D). The hand position at peak acceleration and its variability along the azimuth and depth directions showed no differences across conditions (Figure 5, first column).

**Figure 2:**
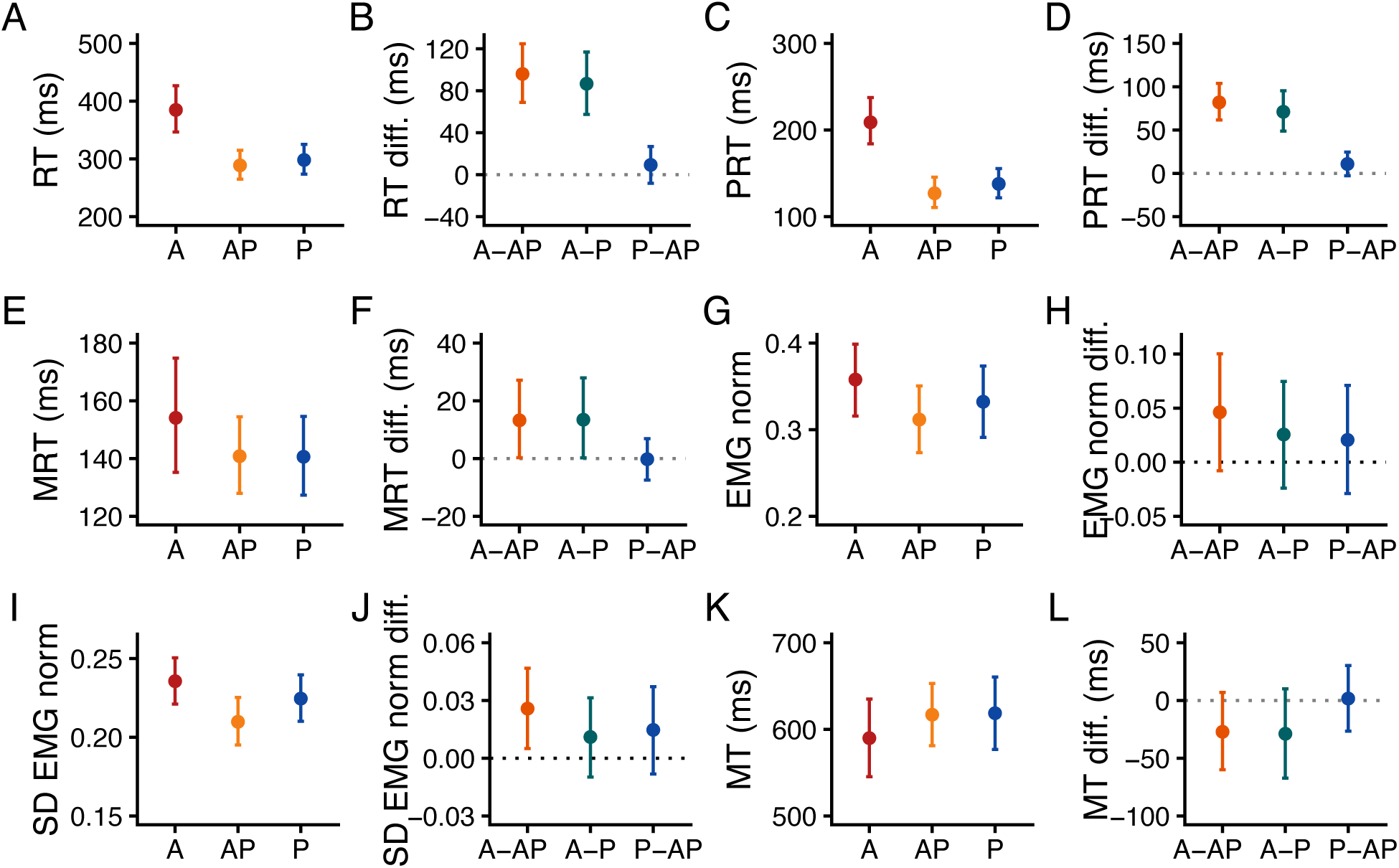
Bayesian model estimates of the Reaction Time (A), Premotor Reaction Time (C), Motor Reaction Time (E) normalized EMG and its variability (G, I) and Movement Time (K). The dots represent the model’s posterior estimates, and the error bars denote the 95% HDI of the estimates. B, D, F, H, J, L) Estimates of the differences in RT, PRT, MRT, normalized EMG, normalized EMG variability, and MT between conditions. The dots represent the estimated difference, and the error bars denote the 95% HDI of the estimates. The grey dotted line represents the points of equality between the compared conditions.

**Figure 3:**
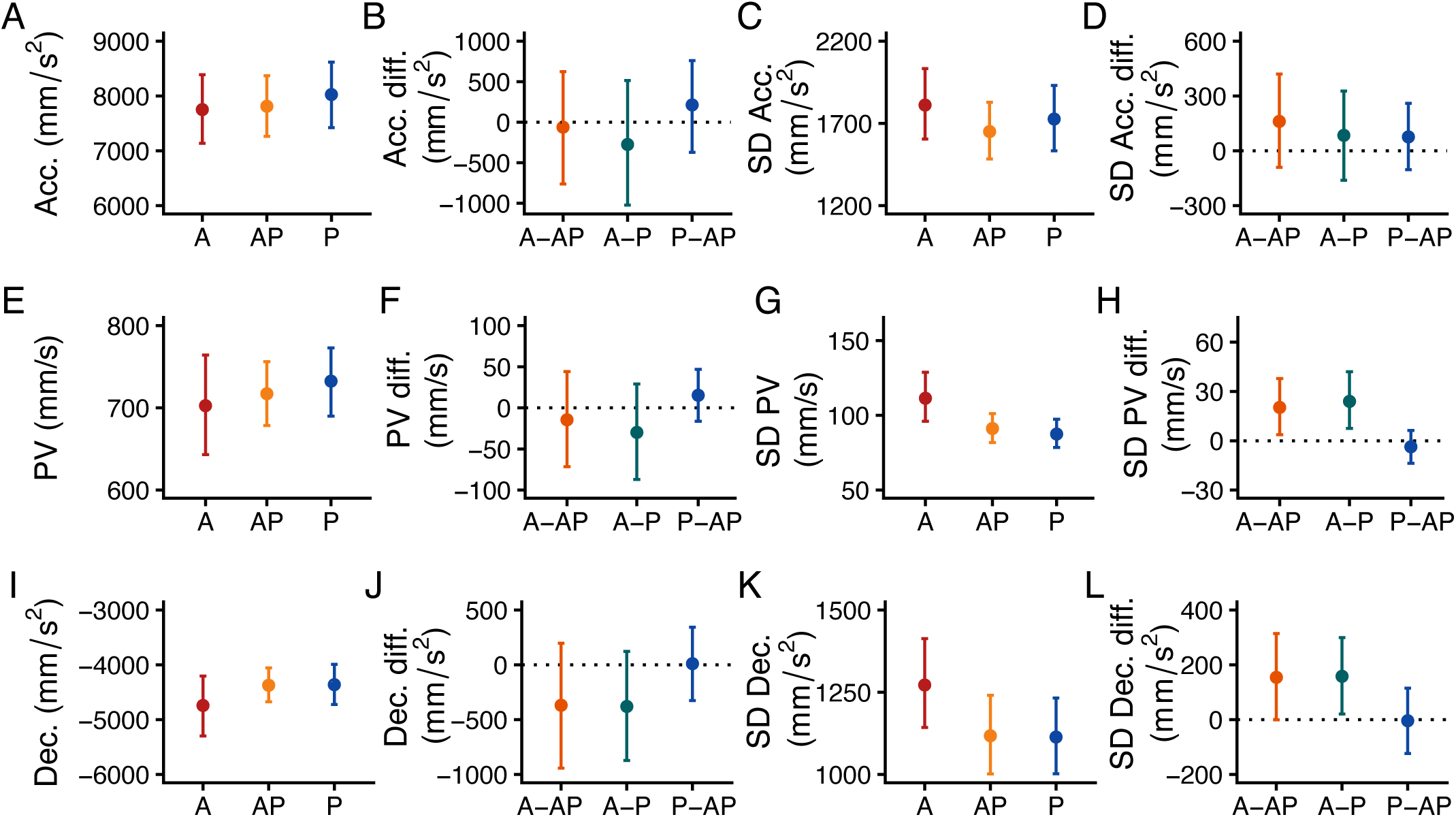
Bayesian distributional regression model estimates of the Acceleration (A), Peak Velocity (E), and De-celeration (I). C, G, K) Estimates of Acceleration, Peak Velocity, and Deceleration variability. The dots represent the model’s posterior estimates, and the error bars denote the 95% HDI of the estimates. B, F, J) Estimates of the differences in Acceleration, Peak Velocity, and Deceleration between conditions. D, H, L) Estimates of the differences in Acceleration, Peak Velocity, and Deceleration variability between conditions. The dots represent the estimated difference, and the error bars denote the 95% HDI of the estimates. The grey dotted line represents the points of equality between the compared conditions.

**Figure 4:**
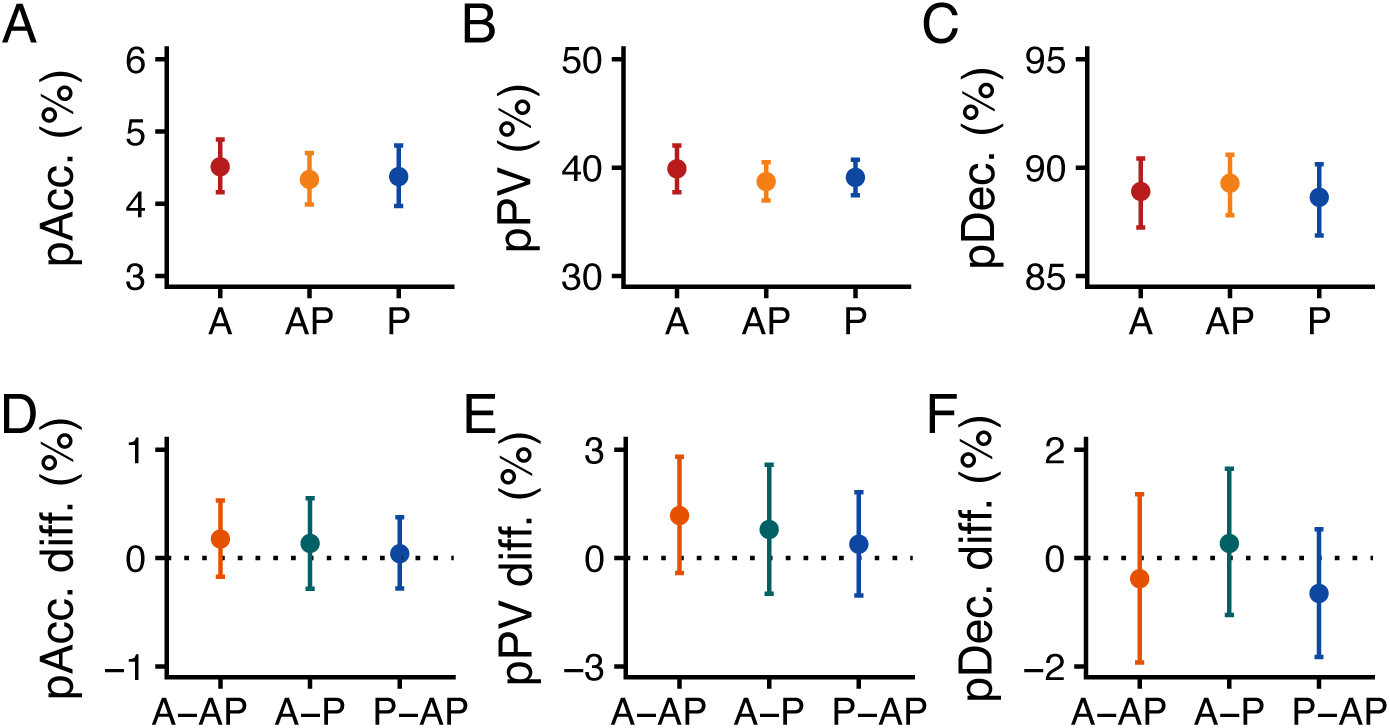
Bayesian regression model estimates of the index finger position at peak acceleration pACC (A), peak velocity pPV (B), and deceleration pDec (C). The dots represent the model’s posterior estimates, and the error bars denote the 95% HDI of the estimates. D, E, F) Estimates of the differences in pAcc, pPV, and pDec between conditions. The dots represent the estimated difference, and the error bars denote the 95% HDI of the estimates. The grey dotted line represents the points of equality between the compared conditions.

**Figure 5:**
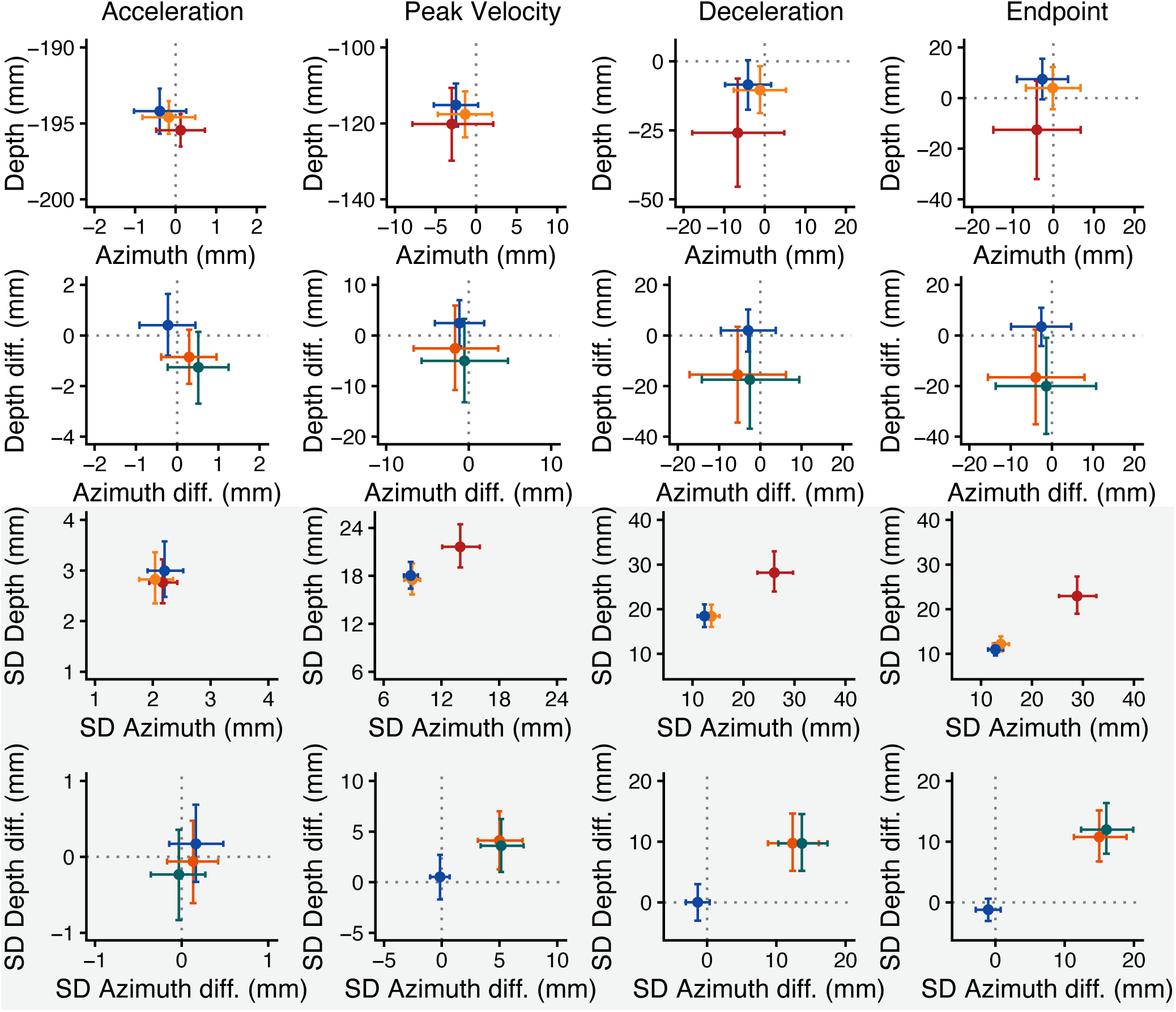
Estimates of the finger distance from the target at Acceleration, Peak Velocity, Deceleration, and Endpoint. Each column represents the results at each kinematic marker. The first row represents the estimated average distance from the target, the second row the estimated differences between conditions, the third row the estimated variability, and the fourth row the estimated differences in variability between conditions for each kinematic marker. The light gray area separates the position from the variability results. In the first and third rows, the dots represent the mean distance from the target and its variability, and the error bars denote the 95% HDI of the estimates. The grey dashed line in the first row represents the center of the targets along the azimuth and depth directions. In the second and fourth rows, the dots represent the difference in mean distance from the target and in its variability, and the error bars denote the 95% HDI of the estimates. The grey dashed line in these rows represents the points of equal position (second row) and equal variability (fourth row) in the depth and azimuth dimensions.

#### Effect of the target position

In general, the effect of the target positions was similar across conditions for all the considered variables. The back-transformed posterior slope estimates for the RT, PRT, and MRT showed a positive but minimal effect of the target position, which corresponds to an average change of approximately ∼4 ms in RT, PRT, and MRT from the front (0 degrees) to the most leftward target position (90 degrees). This effect was similar between conditions. The slopes for the mean normalized EMG were close to 0 at each condition, ranging from 0.0005 to - 0.0005, with no difference between conditions. This corresponds to an average change of approximately ± 0.045% of normalized EMG from the front (0 degrees) to the most leftward position (90 degrees), indicating almost no change of EMG activation across different target positions. Likewise, the SD of EMG did not change across target positions (all slopes equal to 0) and this effect was similar across conditions. The average slope of the Acc was ∼18 mm/s ^2^, and showed no credible differences across conditions. The pAcc did not change with the target positions (slope = 0); such invariance was similar across conditions as well. The hand distance from the target at the peak acceleration was slightly affected by the target position in all conditions, showing an average change of ∼ -0.04 mm in the azimuth and ∼ 0.02 mm in the depth direction. This corresponded to a change of 3.6 mm in azimuth and 1.8 mm along the depth dimension from the front (0 degrees) to the most leftward (90 degrees) target. Such a change was similar across conditions. The hand position variability did not change across target positions for all the conditions.

Taken together, these results indicate a general difference in movement preparation when reaching auditory and proprioceptive or audio-proprioceptive targets. Interestingly, the effect of sensory input was the strongest during the sensory encoding phase (PRT), with almost no effect during the coordination of motor commands (MRT). The reduced effect of the target modality extended to the initial part of the movement, immediately after its onset (i.e., peak acceleration). These findings suggest that, during the coordination of motor commands and movement initiation, actions are partially unaffected by the target modality.

### Movement Execution

#### Effect of the sensory condition

The analysis revealed no differences across conditions for MT, PV, and Dec (Figure 2K, L; Figure 3E, F, I, J). However, we found a slightly higher variability in A compared to P and AP, and a similar variability between P and AP for the PV (Figure 3G, H). We found a similar pattern of variability results for Dec, which, however, was almost not credibly different across conditions (Figure 3K, L). The pPV and pDec were similar across conditions, with pPV occurring at ∼40 %, and pDec at ∼90 % of the movement trajectory (Figure 4). Interestingly, while the index finger distance from the target in azimuth and depth dimensions was similar across conditions at the PV, it was slightly farther from the target at the Dec and the Endpoint for the A compared to the P and AP conditions (Figure 5, second, third, and fourth columns, first and second rows). Moreover, we found a higher movement variability in A compared to P and AP, with a similar variability between P and AP at all the considered positions (Figure 5, second, third, and fourth columns, third and fourth rows).

#### Effect of the target position

In general, the effect of target positions was similar across conditions for all the considered variables. The slopes of PV and Dec, were, respectively, negative (∼ -1.6 mm/s for the PV) and positive (∼ 6 mm/s^2^ for the Dec), and did not change across conditions. The pPV was unaffected by the target position and was similar across conditions. The pDec showed an average change according to the target position of ∼0.004% and was similar across conditions. This corresponds to an average change of 0.34% from the front (0 degrees) to the most leftward position (90 degrees), indicating almost no change of pDec across different target positions. In general, the distance from the target in azimuth and depth at the PV, Dec and Endpoint, and its variability were only minimally (if not) affected by the target position. The finger distance from the target at PV slightly changed according to the target location along the azimuth direction only in the P condition (mean = -0.1 mm, HDI = -0.17 mm, -0.03 mm) and showed an average slope of ∼ 0.09 mm along the depth dimension for all the conditions. The effect of the target position was similar across conditions in both azimuth and depth dimensions. The variability of the finger position in PV changed across target positions in all the conditions along the azimuth direction, with a slope ranging from 0.07 mm in A to 0.02 mm in AP conditions. The variability along the depth was unaffected by the target position. The slope was credibly higher in A compared to AP (mean = 0.05 mm, HDI = 0.01, 0.1) only for the azimuth direction. No other differences were detected across conditions in both directions. The finger position at Dec showed no change in slope in either direction, except in A for the depth dimension (mean = -0.25 mm, HDI = -0.46 mm, -0.04 mm). However, it was similar across conditions. The variability showed an average slope of -0.065 mm along the depth dimension, which was, again, similar across conditions. Finally, the slope at the endpoint showed an average decrease in endpoint error only along the depth dimension of ∼ -0.2 mm, and was similar across conditions. The endpoint variability showed a negative slope in the A condition along the depth dimension (mean = -0.09, HDI = -0.17, 0.02), which was credibly lower than P and AP (average difference = -0.09 mm, HDI -0.17, -0.02 for both comparisons).

Taken together, these results suggest that the hand’s position relative to the target is the movement parameter most strongly influenced by the target-related sensory modality during action execution. In contrast, movement timing, velocity, and deceleration remain largely unchanged across conditions. This suggests that target-related sensory information is mainly used to specify the hand’s position with respect to the sensed target, and that this contribution emerges early in the movement execution, at the peak velocity.

## Discussion

This study aimed to investigate the sensorimotor integration process of non-visual target-related sensory information within the movement preparation and movement execution phases of a reaching movement. To this aim, we asked participants to reach auditory, proprioceptive, and audio-proprioceptive targets while recording their movement kinematics and muscle activation through EMG. Several variables were analyzed. Movement preparation was assessed through analyzing RT, PRT, MRT, EMG activity at MRT, Acc, and the index finger position and variability at Acc. While RT provided general information about the effect of target-related modalities in movement preparation, the analysis of PRT and MRT allowed us to investigate the specific contribution of each sensory modality within the sensory encoding (PRT) and coordination of motor commands (MRT). Movement execution was instead investigated through the analysis of PV, Dec, MT, pAcc, pPV, and pDec, and the index finger position in azimuth and depth at PV, Dec, and Endpoint.

Results showed a modulation of RT according to the target-related sensory modality, evidencing their crucial role during the movement preparation phase. Interestingly, within the RT, the sensory encoding phase was modulated according to the available sensory information, with longer PRT in the auditory than in proprioceptive and audio-proprioceptive conditions, which were similar. In contrast, the coordination of motor commands (MRT) and the EMG activity at MRT were almost unaffected by the target-related sensory information. Our results align with previous EEG findings showing different sensory-related preparatory activity for non-visual targets (Bianco et al, 2020a). Concurrently, MRT and the EMG activity results corroborate studies showing a similar pattern of activation across motor cortices following auditory and somatosensory target presentations (Bianco et al, 2020a; Fiorini et al, 2021; Lucia et al, 2023). These results suggest that target-related sensory information primarily affects the sensory encoding phase rather than the coordination of motor command during the motor preparation. The analysis of the PV, Dec, MT, pAcc, pPV, and pDec showed that, once started, actions were similar in terms of average speed, deceleration, and time to complete the movement. However, confirming and extending our previous study’s results (Camponogara, 2025), the analysis of the distance from the target in azimuth and depth at PV, Dec, and endpoint revealed an effect of target-related sensory information on action variability, showing a higher variability in A compared to P and AP since the early stages of the action (PV). Thus, our results suggest that actions toward auditory and proprioceptive targets may be characterized by a similar modulation of kinematic movement parameters during their execution. However, the target-related sensory modality may primarily be used to define the hand position relative to the target, which changes continuously throughout action execution and reaches its final configuration during the deceleration phase.

The distinct effects of sensory modalities during sensory encoding and the similarities across conditions during the coordination of motor commands and movement initiation suggest a different use of sensory information during action preparation and movement initiation. Previous studies have shown a functional independence between these two phases (Haith et al, 2016). Concurrently, neurophysiological studies on monkeys have shown a change in the pattern of neural activation when transiting from the movement preparation to movement initiation, further confirming their functional separation (Kaufman et al, 2016). In different sensory contexts, studies on fMRI in humans have shown a modulation of brain spatial pattern activity in early action preparation according to the type of sensory modality (vision and audition) (Kreter et al, 2025). Studies analyzing the time dynamics of EEG beta-band activity (the brain’s signature of movement preparation) before and after the movement onset in humans further confirmed these findings, showing a modulation of beta-band before the movement onset according to the target uncertainty (Tzagarakis et al, 2010). It is thus plausible to think that within the movement preparation stage, two processes occur: one related to the elaboration of the target information and deciding “when” to act, and one related to defining a general movement coordination pattern to start the action (Kreter et al, 2025).

In this regard, the transition from the sensory encoding to the coordination of motor commands, and the transition from this latter to the movement onset may be regulated by a double-threshold accumulation gated mechanism (Servant et al, 2021; Dendauw et al, 2024). This model posits that each transition between stages in motor preparation is triggered once a specific accumulation threshold is reached. During the sensory encoding phase, the nervous system accumulates information about the target’s location over time. A first gating threshold determines the level of sensory evidence accumulation to initiate the motor coordination phase. The rate of this accumulation depends on the quality of the sensory input: when the available sensory information is more uncertain, the accumulation proceeds more slowly; conversely, more reliable information enables faster accumulation (Servant et al, 2021). According to our results, the lower uncertainty characterizing proprioception (Camponogara, 2025) may facilitate the sensory accumulation process, allowing for a faster transition from the sensory encoding (i.e., PRT) to the coordination of motor commands (i.e., MRT) than in the auditory condition. The coordination of motor commands, instead, reflects the accumulation of neural drive to the muscles, governed by a second gating threshold. Once this second threshold is exceeded, movement begins, and changes in kinematics become observable (Servant et al, 2021). Again, according to our results, this phase appears to be relatively unaffected by the incoming sensory information about the target, and most of the sensorimotor processes occurring in this phase may be mainly related to the coordination of motor commands. However, further studies are needed to fully understand this process under different sensory conditions.

The similarities across non-visual sensory modalities during movement initiation contrast with previous results on visual/proprioceptive target reaching, which often showed an effect of the target modality since the movement initiation phase (Sarlegna and Sainburg, 2007; Sober and Sabes, 2005; Sarlegna et al, 2009). A plausible explanation for such a discrepancy may lie in the sensory modalities involved in sensing the *reaching hand* and the *target* in visual vs. proprioceptive and auditory vs. proprioceptive target reaching. In visual target reaching, the *reaching hand* position is defined through multisensory inputs (vision and proprioception), whereas the *target* is defined through unisensory input (vision). In contrast, in proprioceptive target reaching, both the *reaching hand* and the *target* are defined within the same sensory modality. Thus, the different movement preparation in visual vs. proprioceptive target reaching may largely reflect the availability of multisensory or unisensory information of the *reaching hand*, as the former usually leads to a better action performance than the latter (Camponogara, 2023). This has been further confirmed by studies showing that a modulation of the *reaching hand* visual information impacts movement preparation toward a visual target (Sober and Sabes, 2005; Sarlegna and Sainburg, 2007; Sarlegna et al, 2009). In auditory vs. proprioceptive target reaching, the position of the *reaching hand* is defined solely by proprioceptive information, whereas the *target* is sensed through two different modalities: proprioception or audition. Consequently, while in proprioceptive condition, the *reaching hand* and the *target* position are monitored within the same sensory modality; in the auditory condition, they are defined through two different sensory modalities, thus requiring a cross-modal comparison (Camponogara, 2025). Our results suggest that differences in the *reaching hand* and the *target* comparisons across conditions primarily affect the sensory encoding phase of movement preparation and action execution. However, it does not dramatically affect the coordination of motor commands and the movement initiation phase.

The observed sensorimotor temporal dynamics align with Woodworth’s classic two-stage model of movement control in visual target reaching (Woodworth, 1899). This model posits an initial impulse phase, corresponding to early movement execution and reflecting motor preparation processes, followed by a limb-target control phase, during which online sensory information is used to refine and guide the movement (for review, see Elliott et al (2010)). Our results extend these findings in the non-visual (auditory and proprioceptive) domain by demonstrating that in the initial impulse phase (i.e., the movement onset), non-visual target-related sensory inputs have a lower impact than in the limb-target control phase (i.e., early and final movement execution), which, instead, modulates according to the target-related sensory information. Taken together, our results suggest that reaching performance toward non-visual targets is governed by four distinct phases (Figure 6). The first involves the sensory encoding, which is influenced by the available target-related sensory modalities. The second phase corresponds to the coordination of motor commands, which are relatively unaffected by the incoming target-related sensory information and trigger the initiation of movement. A third phase is the initial movement onset, which marks the transition from the movement preparation to the movement execution. This phase is independent of the available target-related sensory information. Lastly, the fourth phase is characterized by a shift from the initial movement onset to the early and late execution, during which available sensory information is integrated to refine and adjust the movement trajectory.

**Figure 6:**
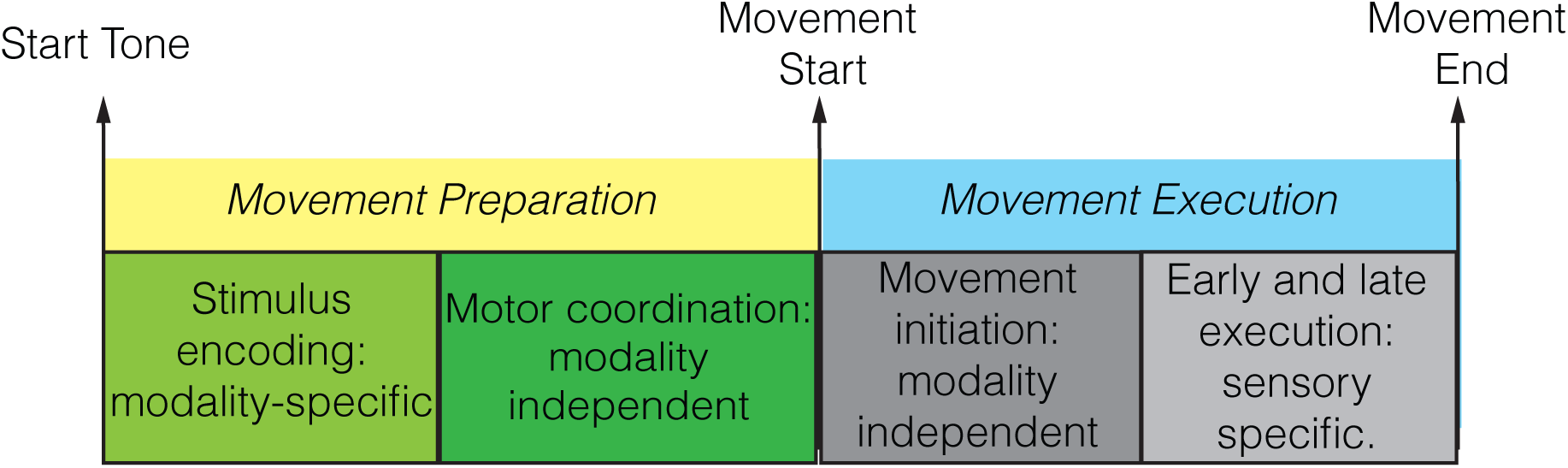
Representation of the four phases of movement control. Target-related sensory information is mostly relevant during the stimulus encoding phase of movement preparation and early/late stages of movement execution. In contrast, it is less relevant during the motor coordination phase of movement preparation and the initial movement execution.

These findings provide further support for the sensorimotor integration framework proposed by Camponogara (2023) for visual and proprioceptive target reaching, extending it to auditory and proprioceptive modalities. It is well established that, during movement preparation, sensory estimates of the future state of the reaching hand are integrated into the motor plan to generate predictive representations of the forthcoming movement. During movement execution, these predictions are refined through the integration of incoming sensory inputs (Diedrichsen et al, 2010; Brenner and Smeets, 2025). According to the model proposed by Camponogara (2023), motor preparation involves the integration of proprioceptive (or haptic) estimates of the reaching hand with the available target-related sensory information (Friston et al, 2011; Sandbrink et al, 2023; Camponogara, 2023, 2025). At the movement onset, the release of motor commands is primarily guided by proprioceptive (or haptic) estimations of the future state of the reaching hand. As movement unfolds, these predictions are continuously updated with real-time sensory information from both the reaching hand and the target (Camponogara, 2023). In the context of non-visual target reaching, our results suggest that during the sensory encoding phase of movement preparation, proprioceptive information from the *reaching hand* and the *target* -related sensory information (either proprioceptive or auditory) are accumulated to generate sensorimotor predictions about the upcoming movement. Once enough sensory information is accumulated, the transition to the coordination of motor commands occurs, initiating the movement (Servant et al, 2021; Dendauw et al, 2024). In *non-visual* target reaching, this phase is relatively unaffected by the *target* -related sensory information and may be primarily controlled by sensorimotor predictions (Harris and Wolpert, 1998). As movement progresses, the sensorimotor predictions are dynamically adjusted by integrating upcoming sensory information from the *reaching hand* (proprioception) and the *target*, enabling fine-tuned control of the action. Movement execution is strongly affected by the target-related sensory modality, impacting the trial-to-trial movement variability.

In conclusion, our results indicate that actions directed toward non-visual targets are characterized by a modulation of the sensory encoding, but not the coordination of motor commands during movement preparation. Concurrently, while the initial movement execution is unaffected by the target-related sensory modality, early and late stages of movement execution are modulated according to the available sensory information. These findings suggest that, in auditory and proprioceptive target reaching, target-related sensory inputs mainly affect the sensory encoding of movement preparation and the control of action performance during early and late phases of movement execution. In contrast, sensory information plays a limited role during the coordination of motor commands and in the movement initiation. These results open new perspectives in sensorimotor control.

The vast majority of studies comparing *visual* and *non-visual* (proprioceptive) targets often reported a differ-ence in both movement preparation and action execution. The similarities in specific phases of the action when interacting with pure *non-visual* targets (auditory and proprioceptive) we found in this study suggest that, when interacting with auditory or haptic targets, motor preparation and execution may adhere to alternative sensorimotor control principles than those outlined in visuo-proprioceptive studies.

## CRediT author statement

**Ivan Camponogara:** Conceptualization, Methodology, Software, Validation, Formal analysis, Investigation, Data Curation, Writing - Original Draft, Writing - Review and Editing, Visualization, Supervision, Funding acquisition.

## Acknowledgements

We acknowledge the support of the Zayed University Research Incentive Fund (grant 23076). We thank Prof. Dr. Jeroen Smeets for his insightful comments on an earlier version of this manuscript.

## Additional information

### Competing interests

The authors declare no competing interests.

Correspondence and requests for materials should be addressed to I.C.

